# Bayesian optimization for demographic inference

**DOI:** 10.1101/2022.09.06.506809

**Authors:** Ekaterina Noskova, Viacheslav Borovitskiy

## Abstract

**Motivation:** Inference of demographic histories of species and populations is one of the central problems in population genetics. It is usually stated as an optimization problem: find a model’s parameters that maximize a certain log-likelihood. This log-likelihood is often expensive to evaluate in terms of time and hardware resources, critically more so for larger population counts. Although genetic algorithm based solution have proven efficient for demographic inference in the past, it struggles to deal with log-likelihoods in the setting of more than three populations. Different tools are therefore needed to handle such scenarios.

**Results:** We introduce a new specialized optimization pipeline for demographic inference with time-consuming log-likelihood evaluations. It is based on Bayesian optimization, a prominent technique for optimizing expensive black box functions. Comparing to the existing widely used genetic algorithm solution, we demonstrate new pipeline’s superiority in time limited conditions for demographic inference with four and five populations when using log-likelihoods provided by the *moments* tool. Moreover, we expect this behavior to generalize just as well to other expensive-to-evaluate log-likelihood functions in the field.

**Availability:** The proposed method was implemented as part of the *GADMA* software framework and is freely and openly available on GitHub: https://github.com/ctlab/GADMA.

**Supplementary information:** Supplementary materials are available as a separate document.

## 1 Introduction

The history of populations’ development — the *demographic history* — is imprinted into genomes of all individuals. It is a record of events that happened in the past, including changes in population size, population splits, migration and selection events. Demographic histories provide important insights about populations to biological and medical researches.

Reconstruction of the demographic history is called *demographic inference*. There exists a multitude of tools for demographic inference from genetic data based on a variety of mathematical models (Gutenkunst *et al*., 2009; Jouganous *et al*., 2017; Steinrücken *et al*., 2019; Kamm *et al*., 2020; Excoffier *et al*., 2013, 2021; DeWitt *et al*., 2021). They all consist of two rather independent components: *simulation* and *optimization*. The simulation component evaluates *log-likelihood* of the observed data under a proposed demographic history. The optimization component takes in a *demographic model* — a parametric family of demographic histories — and searches for the parameters that maximize log-likelihood produced by the simulation component.

Existing tools for demographic inference are usually limited in the number of analyzed populations. For example, the original version of the *∂*a*∂*i tool by Gutenkunst *et al*. (2009) is able to handle only up to three populations. Another tool, *moments* by Jouganous *et al*. (2017) can handle only up to five populations. This is caused by computational complexity of simulation techniques used therein, which would scale exponentially with the number of populations: for example, *∂*a*∂*i numerically solves a partial differential equation whose dimension is equal to the number of populations. Some tools, e.g. *momi2* by Kamm *et al*. (2020), use methods that scale linearly with respect to the number of populations, and are therefore able to handle an arbitrary number thereof. This does not change the big picture though: different tools rely on different mathematical models with different assumptions and thus are not interchangeable.

In the end, demographic inference for multiple populations is widely regarded to be a slow and expensive procedure, making the problem of speeding it up to be a valuable research direction. One way to do this, is to speed up the simulation component of the tools. Along these lines *moments* was presented as a faster but somewhat less accurate alternative to *∂*a*∂*i as well as *∂*a*∂*i itself received GPU support in Gutenkunst (2021) that allowed it to handle up to five populations. A whole other direction to boost demographic inference is to improve the optimization component. This was done in the first author’s previous work. It proposed a new tool *GADMA* (Noskova *et al*., 2020, 2022) based on a genetic algorithm as a substitute to the optimization components of various tools — mostly based on the (restarted) local search techniques — giving better performance.

However, even though e.g. *∂*a*∂*i’s and *moments*’s simulation components can now handle up to five populations, it is recommended to use *GADMA* with up to three populations. This is a natural limitation stemming from genetic algorithm’s hunger for log-likelihood evaluations, something that becomes prohibiting for really expensive log-likelihoods.

In this paper we address this problem by introducing *Bayesian optimization* as a tool for demographic inference with larger numbers of populations and implementing it as part of the *GADMA* framework. Bayesian optimization (Shahriari *et al*., 2015) is a state of the art approach for optimizing expensive-to-evaluate functions within a limited (time) budget. From the data acquired by evaluating the target function it *learns* a probabilistic model and uses it to guide the choice of a new evaluation location. It comes with a cost though: Bayesian optimization’s inner workings are rather expensive, making it suitable only for problems where the target function is itself expensive to evaluate. Previous works used it for tuning large-scale systems (Snoek *et al*., 2012; Chen *et al*., 2018) or robot control policies (Berkenkamp *et al*., 2021; Jaquier *et al*., 2022) as well as for chemical reaction optimization (Shields *et al*., 2021), to name a few.

Our contribution is the following. We propose and implement a specialized Bayesian optimization pipeline for demographic inference with larger population counts. To choose this pipeline, we evaluate the performance of a number of candidates using *moments*’s simulation component. In the same way we then study the efficiency of the approach, expecting it to generalize to other expensive-to-evaluate settings. The results show rapid convergence of the proposed method on different datasets and prove its efficiency compared to the genetic algorithm in the settings of four and five populations.

## 2 Methods and materials

In this section we introduce the methods used in this study, namely Bayesian optimization and Gaussian process regression, the machine learning technique Bayesian optimization relies upon. In the end, we describe the datasets used in Section 4 for performance evaluation.

### 2.1 Bayesian optimization

As it was mentioned in the introduction, *Bayesian optimization* provides state of the art performance for optimizing expensive black-box functions.

Bayesian optimization minimizes a *black box function ϕ* : 𝒯→ ℝ, i.e. a function we are able to evaluate at any input *t* ∈ 𝒯, obtaining a possibly noisy observation *y*(*t*) of *ϕ*(*t*), but nothing more (e.g. no gradients). Moreover, it is usually assumed that each evaluation is expensive and the objective is to converge as fast as possible or to get as close to the global optimum as possible within a fixed budget of evaluations or time.

The main idea of this technique is to use a relatively cheap surrogate model to approximate the expensive target function *ϕ* each iteration and use it as a proxy to guide decisions. The most widely used surrogate models are Gaussian processes (Rasmussen and Williams, 2006) discussed in detail in Section 2.2. The reason is their ability to perform well in small data regimes and to quantify uncertainty associated with their own predictions.

The optimization procedure begins by drawing a small random sample from the domain 𝒯 and evaluating the target function at each of the inputs from this sample. The obtained data is called the *initial design*. After this, a prior Gaussian process is chosen, usually from the options detailed later in Section 2.2.4. This may be done manually, aided by some external considerations like prior knowledge about smoothness of the target function, or alternatively it may be chosen by means of the cross-validation procedure described in Section 2.2.5.

At each optimization iteration, using as data the target function evaluations *t*_1_, *y*_1_, …, *t*_*n*_, *y*_*n*_ obtained so far, Gaussian process regression is executed, as detailed in Section 2.2, resulting in a posterior Gaussian process 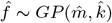. The value 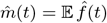 of its mean function 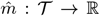 at input *t* is treated as a *prediction* of the target function therein, while the value 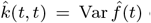 of its covariance function 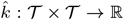 at (*t, t*) is treated as the projected variance of this prediction, i.e. a *measure of uncertainty*.

Given the posterior Gaussian process, the location *t*_∗_ to evaluate the target function next is chosen by solving

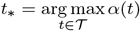

where *α* : 𝒯 →ℝ is the *acquisition function* defined in terms of the posterior process 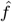 (Frazier, 2018). Common choices of *α* include

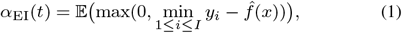

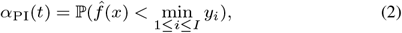

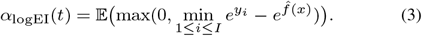

Here “EI” stands for Expected Improvement and “PI” stands for Probability of Improvement. The acquisition function *α*_logEI_(*t*) by Hutter *et al*. (2009) is intended for use in conjunction with log-transformed data, i.e. when *y*_*j*_ *>* 0 and *t*_1_, log *y*_1_, …, *t*_*n*_, log *y*_*n*_ are fed into Gaussian process regression instead of *t*_*i*_, *y*_*i*_, …, *t*_*n*_, *y*_*n*_.^1^ For a Gaussian process 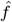 with known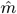 and 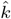 these acquisition functions are tractable and may be computed in closed form and efficiently optimized by gradient descent (with restarts). Further we refer to the acquisition functions in Equations (1), (2) and (3) by the names `EI`, `PI` and `LogEI` respectively.

A part of Bayesian optimization’s workflow is illustrated in Figure 1.

**Fig. 1:**
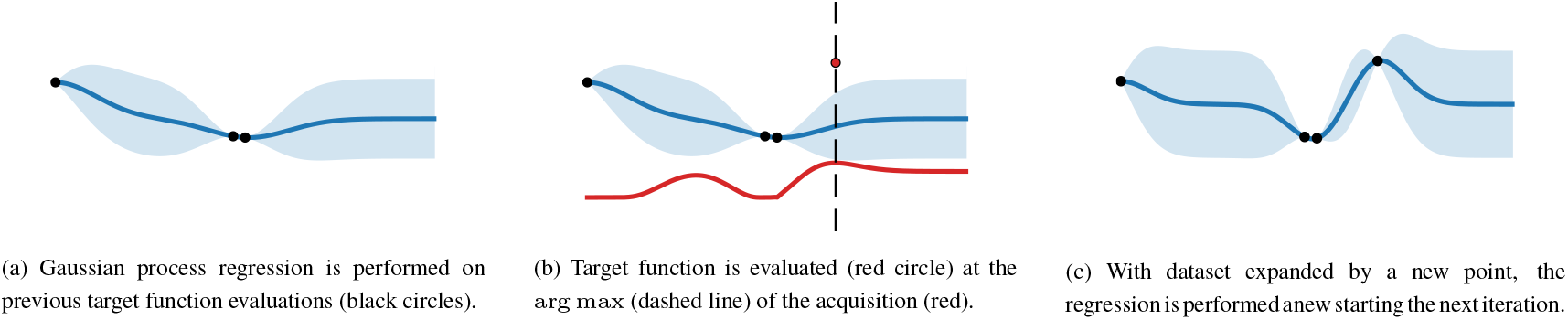
A fragment of Bayesian optimization’s workflow. The blue line is the prediction of the Gaussian process regression, i.e. 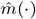, the shaded blue regions represent uncertainty bars whose hight is proportional to. 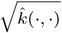. The red curve is the acquisition function, the dashed vertical line is its arg max.

### 2.2 Gaussian process regression

Gaussian process regression models an unknown function from data and provides well-calibrated uncertainty bars alongside with its predictions. As its name suggests, it is based on *Gaussian processes*.

#### 2.2.1 Gaussian processes

Mathematically, a *Gaussian process* is a family {*f*_*t*_}_*t*∈𝒯_ of jointly Gaussian random variables *f*_*t*_ indexed by some set 𝒯. Sometimes people reserve this term only for such index sets 𝒯 that 𝒯 ⊆ ℝ but we will not stick to this convention. Instead, 𝒯 will be a subset of ℝ^*d*^, typically a product of segments, corresponding to the domain of the modeled function.

A Gaussian process (or rather, strictly speaking, its distribution) is determined by a pair of deterministic functions: the *mean function m* : 𝒯 → ℝ and the *covariance kernel k* : 𝒯 × 𝒯 → ℝ:

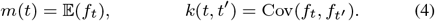

Conversely, each pair of deterministic functions *m* and *k* where *k* is positive semidefinite (see Rasmussen and Williams (2006) for the definition) constitutes a Gaussian process. This one-to-one correspondence motivates the standard notation

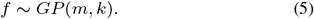

Because of their simplicity, Gaussian processes are attractive to encode distributions over functions. Furthermore, they turn out to be particularly suitable for the *Bayesian learning* framework that we briefly describe next.

#### 2.2.2 Bayesian learning

The framework of *Bayesian learning* combines a *prior* Gaussian process *f* ∼ *GP* (*m, k*), some data ***y*** ∈ ℝ^*n*^ at locations ***t*** ∈ 𝒯^*n*^ and a likelihood function *p*(***y*** | *f*) into the *posterior* process 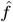 given by the Bayes’ rule^2^

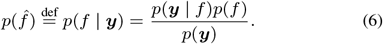

The posterior 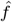 is “similar” to the prior *f* but also respects the data ***y*** in a way prescribed by the likelihood *p*(***y*** | *f*).

The posterior process 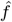 may in general fail to be Gaussian, in which case any computations involving it will require complex numerical techniques like Markov Chain Monte Carlo (MCMC). Fortunately, if the likelihood function corresponds to the situation where for each observation *y*_*i*_ one assumes *y*_*i*_ = *f* (*t*_*i*_) + *ε* with 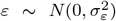 being Gaussian noise with variance 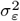, then 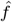 is a Gaussian process. Moreover, 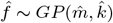 with functions 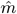 and 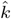 given by the following simple closed form expressions (Rasmussen and Williams, 2006) which we present, for further simplicity, under the assumption that *m* ≡ 0.

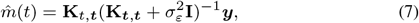

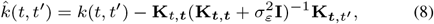

where for a pair of vectors ***a*** ∈ 𝒯^*l*^, ***b*** ∈ 𝒯^*s*^, the symbol **K**_***a***,***b***_ denotes the *l* × *s* matrix defined by (**K**_***a***,***b***_)_*ij*_ = *k*(***a***_*i*_, ***b***_*j*_); ***t*** = (*t*_1_, …, *t*_*n*_)^T^ and similarly ***y*** = (*y*_1_, …, *y*_*n*_)^T^. Because of this remarkable simplcity, such likelihood assumption is often the setting of choice for applications.

Bayesian learning with Gaussian processes is the main part of the *Gaussian process regression* technique which we describe next.

#### 2.2.3 Regression

As a preparatory step before *Gaussian process regression*, a parametric family of *priors f*_*θ*_ ∼ *GP* (*m*_*θ*_, *k*_*θ*_) must be selected. We postpone the discussion on specific families of priors used in practice until Section 2.2.4, but right away make the widely used assumption of *m*_*θ*_ ≡ 0 for simplicity.

After choosing such a family, the Gaussian process regression technique proceeds in two steps. First, the optimal parameters 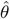 of the prior and 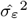 of the likelihood function are found by maximizing the marginal log-likelihood (Rasmussen and Williams, 2006):

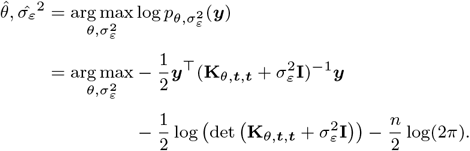

To solve this optimization problem the simple gradient descent with multiple restarts is usually the tool of choice.

The second step of the Gaussian process regression is to compute the Cholesky decomposition of the positive-definite matrix 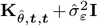, i.e. lay the groundword for solving linear systems of form

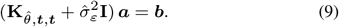

After this is done, the prediction at an arbitrary location *t* is given by 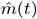 and the error bars therein are given by 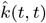.

The optimization problem which involves solving a linear system and computing a determinant, as well as the later step of computing the Cholesky decomposition have computational complexity O(*n*^3^) where *n* is the size of the dataset.^3^ This prohibits the use of this simple approach for large *n*. However, after the two steps are completed, the computation of predictions and error bars requires only the relatively cheap — O(*n*) and O(*n*^2^) respectively — and easily parallelizable matrix-vector products.

We now turn to the question of practical Gaussian process priors.

#### 2.2.4 Practical priors

In order to perform Gaussian process regression, a parametric family of priors *f*_*θ*_ ∼ *GP* (*m*_*θ*_, *k*_*θ*_) has to be selected. In practice, the prior mean *m*_*θ*_ is usually assumed to either be zero or an (optimizable) constant. With that said, we focus on choosing *k*_*θ*_ and assume *m*_*θ*_ ≡ 0.

The most widely used family of prior Gaussian processes is the *Matérn family* (Stein, 2012; Rasmussen and Williams, 2006). It is parameterized by three positive parameters: smoothness *ν* that determines how many derivatives the resulting Gaussian process will have, length scale *κ* which scales the *t* axis and variance *σ*^2^ which sacales the *y* axis. These are the zero-mean Gaussian processes with covariances given by

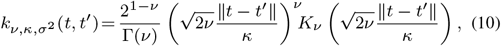

where *K*_*ν*_ is the Bessel function of the second kind and Γ denotes the gamma function (see Gradshteyn and Ryzhik (2014) for definitions).

The general Matérn family given by equation (10) is often divided into subfamilies corresponding to a single value of *ν*. Specifically, the cases of *ν* ∈ {1*/*2, 3*/*2, 5*/*2, ∞} are usually considered, in which Equation (10) may be substantially simplified (Rasmussen and Williams, 2006) to give

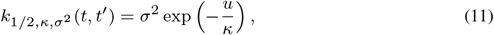

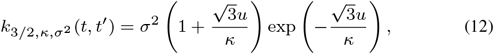

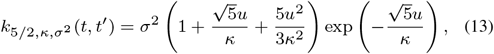

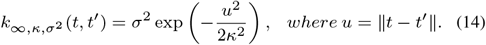

The first covariance is also called the *exponential* kernel, the last — the *Gaussian*, the *squared exponential* or the *RBF* kernel. Subsequently, we refer to zero mean Gaussian processes with kernels (11), (12), (13) and (14) by `Exponential`, `Matern32`, `Matern52` and `RBF` respectively.

#### 2.2.5 Cross-validation for prior selection

Although a parametric family of priors is often chosen manually, there are ways to automate this. Here we describe one popular way, termed *leave-one-out cross validation* (LOO-CV). It takes in some data, usually the initial design, and a finite number of parametric families of priors. For each it computes a certain score, choosing the family with the highest.

Assuming a fixed parametric family of priors, the score is computed by leaving one datum *t*_*i*_, *y*_*i*_ out of the data and computing the prediction quality at *t*_*i*_ of the model 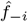 obtained from the rest of the data, which we denote by ***t***_−*i*_, ***y***_−*i*_. This is then averaged over all *i* ∈ {1, …, *n*}:

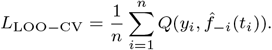

Assuming 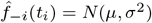, the prediction quality metric *Q* may be defined by

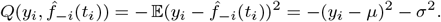

However, this metric clearly favors models that underestimate predictive uncertainty: a lower value of *σ*^2^ directly causes *Q* to be higher. Because of this, another prediction quality metric is used, based on the likelihood, which is able to balance prediction quality with uncertaintly calibration:

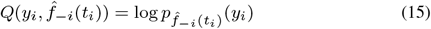

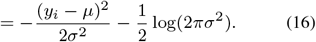

Note that when `LogEI` acquisition function is used, *t*_*i*_, log *y*_*i*_ must be used in Equation (16) in place of *t*_*i*_, *y*_*i*_.

### 2.3 Datasets

The log-likelihood functions provided by simulation components of different tools are determined by the observed genetic data. To evaluate the performance of different optimization pipelines, we use several datasets from the package `deminf_data`.^4^ The genetic data in each of the datasets is represented by the allele frequency spectrum^5^ statistic. Additionally, each dataset contains a demographic model and bounds for the model’s parameters. Datasets are named according to the convention described in Figure 2. More information about the datasets is available in the repository of `deminf_data`. The list of used datasets along with the corresponding times of log-likelihood evaluation is presented in Figure 3.

**Fig. 2:**
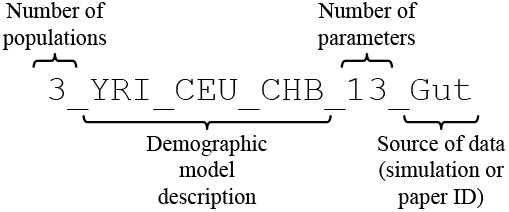
Dataset naming convention.

**Fig. 3:**
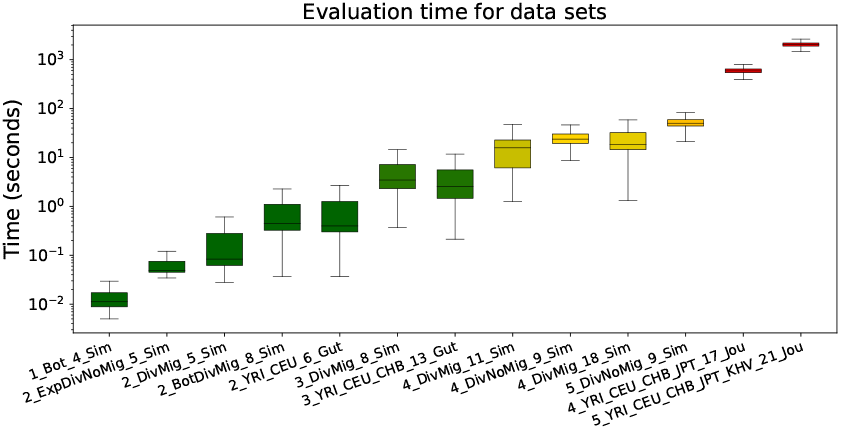
Evaluation times of log-likelihood by the *moments* tool for all used datasets. Y-axis is in log-scale. Boxplots are colored according to median value.

Different Bayesian optimization pipelines studied in Section 4 are tested on the first eleven datasets. There is one dataset with one population, four datasets with two populations, two datasets with three populations, three datasets with four populations and one dataset with five populations.

In Sections 4.6 and 4.7, additionally, two last datasets from Figure 3 are used to (1) compare the final Bayesian optimization pipeline to the genetic algorithm implemented as part of the *GADMA* tool and (2) show that the former is able to find demographic models with higher log-likelihood values than observed in the literature. These datasets correspond to demographic models with four and five modern human populations and the genetic data built by Jouganous *et al*. (2017) from autosomal synonymous sequence data of the publicly available 1000 Genomes Project (Sudmant *et al*., 2015; Consortium *et al*., 2015). The data for four populations includes: 1) Yoruba individuals from Ibadan, Nigeria (YRI); 2) Utah residents with northern and western European ancestry (CEU); 3) Han Chinese from Beijin, China (CHB); and 4) the Japanese from Tokyo (JPT). The data for five populations includes the same four populations and 5) Kinh Vietnamese (KHV) population. These two datasets are also included in `deminf_data` package and are named `4_YRI_CEU_CHB_JPT_17_Jou` for four populations and `5_YRI_CEU_CHB_JPT_KHV_21_Jou` for five populations.

## 3 Implementation

All Bayesian optimization pipelines discussed further below were implemented as part of the open-source software *GADMA* available at https://github.com/ctlab/GADMA.

We highlight several of the ready-to-use libraries implementing Bayesian optimization: GPyOpt (authors, 2016), BOTorch (Balandat *et al*., 2020) and SMAC (Hutter *et al*., 2011; Lindauer *et al*., 2022). Unfortunately, GPyOpt is no longer supported since 2020. BOTorch is actively developed and popular, but SMAC proved itself well in a number of applications (Lago *et al*., 2018; Hewamalage *et al*., 2021; Wu *et al*., 2022) and, importantly, supports the `LogEI` acquisition function (Hutter *et al*., 2009) that will turn out to be a part of the best performing pipeline later in the following section. Because of this, we use SMAC v0.13.1 to implement Bayesian optimization pipelines within *GADMA*.

## 4 Approach and results

In this section we propose and compare different candidate Bayesian optimization pipelines in the setting of demographic inference. After choosing the most fitting pipeline we compare its performance to the genetic algorithm implemented as part of *GADMA* and show that it is able to attain unmatched log-likelihood values in one real data setting.

### 4.1 Overview

To evaluate the performance of different candidate pipelines we analyze convergence plots compiling 64 independent optimization runs using the datasets from Section 2.3 and the *moments*’ simulation engine. We use *moments* because, while being one of the most popular demographic inference tools alongside with *∂*a*∂*i, it is not so time consuming as *∂*a*∂*i, making it easier to do experimental evaluation. For the latter we consistenly use the same hardware (Intel® Xeon® Gold 6248).

Candidate pipelines are determined by the choice of prior and acquisition function. We consider four zero mean Gaussian process priors with kernels from Equations (11)–(14) and three acquisition functions given by Equations (1)–(3), meaning that overall there are 12 candidates. We denote the candidates by `Acquisition + Kernel`, for example `LogEI + Matern32`. Initial design is always taken to be of size 2*d* where *d* is the number of demographic model’s parameters. It consists of pairs *t*_*i*_, *y*_*i*_ with input locations *t*_*i*_ sampled randomly from the same distribution that *GADMA* uses for its genetic algorithm implementation.

First, we appeal to choosing the prior by comparing the leave-one-out cross validation scores as it was described in Section 2.2.5. We use a large collection (2,000) of points and evaluate the scores for 11 datasets from Section 2.3. The setup and results are detailed in Section 4.2.

After this, we perform the most natural evaluation of the candidate pipelines given the setting: we compare their convergence performance on each of the datasets. This experiment is detailed in Section 4.3. We observe `Matern52` and `RBF` kernels with `PI` and `LogEI` acquisition functions to show more or less equally good results, outperforming other candidates.

After this, we consider cross validation scores on initial design as a tool for automatic prior selection, introducing additional candidate pipelines denoted by `Acquisition + Auto`. We evaluated performance of these for two acquisition functions that performed best on the previous step. Both show equally good or better results compared to the four best performers of the previous step. The details may be found in Section 4.4

Finally, we propose an ensemble pipeline denoted by `Ensemble`. On each optimization iteration a coin is flipped. On heads the `PI` acquisition is used, on tails — `LogEI`. The prior is chosen by comparing the cross validation scores, but in this case only between `Matern52` and `RBF` which performed best in the previous experiments. This last candidate shows better or equal performance compared to the previous champions and is chosen as the final pipeline. See the detailed results in Section 4.5.

To conclude, we compare the final pipeline `Ensemble` to *GADMA*’s genetic algorithm in terms of convergence speed as per iteration and as per wall clock time, showing promising results that are presented in Section 4.6. Immediately after, in Section 4.7, we show that `Ensemble` can give better log-likelihood values then reported so far in the literature for a real data case study.

### 4.2 Comparing cross validation scores to choose a prior

One popular way of selecting the prior is by comparing the leave-one-out cross validation scores (LOO-CV) as described by Section 2.2.5. In an attempt to do so, we evaluated LOO-CV on 2,000 points in 11 datasets from Section 2.3 and used those to compare four prior Gaussian processes under consideration. For each dataset, evaluation points were generated not-uniformly: the first 1,000 points were generated uniformly within target function’s prescribed domain, but the second 1,000 points were generated using the distribution used in initial design procedure. Since the acquisition `LogEI` runs Gaussian process regression on the log transformed data, we also evaluate the LOO-CV scores in this setting.

The results presened in Table S1 and S2 suggest that `Exponential` usually has worst performance and `Matern52` is most often the best. However, there were outliers, for example the LOO-CV score for `4_DivMig_18_Sim` dataset was best for `Exponential`. Overall, there is no single clear champion among the priors in terms of the LOO-CV score, however `Matern52` comes closest.

### 4.3 Evaluating 12 basic candidates

The most comprehensive — although not very formal — way to evaluate the performance of different optimization pipelines is by examining the convergence plots, like the one given in Figure 4. The somewhat informal criteria of comparison are: speed of convergence, interquartile distance and, most importantly, the end result. Convergence plots for all datasets and optimization running for 200 iterations can be found in Figures S1–S3.

**Fig. 4:**
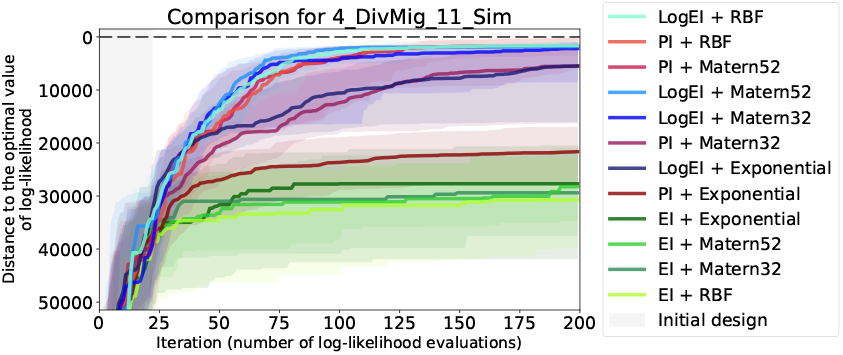
Convergence plot of 12 basic Bayesian optimization pipelines for the `4_DivMig_11_Sim` dataset. For each candidate pipeline 64 optimization runs were independently performed. Solid lines of different colors visualize the median of the sample of 64 values on each iteration, while shaded regions visualize ranges between the first and third quartiles. Grey area indicates the initial design, where random search is performed. The labels in the legend are sorted according to the final median.

Candidates with `Exponential` prior or `EI` acquisition showed poor convergence for all of the 11 datasets. `Matern32` usually showed equal or worse performance compared to `Matern52` (see e.g. Figure 4). As the result, 4 candidates that combine `Matern52` and `RBF` priors with `PI` and `LogEI` acquisitions were considered the most fitting to our setting.

We should note that results of examining the convergence plots are not very well aligned with the cross-validation scores discussed above. They agree in identifying `Exponential` as the worst choice, but disagree in some other cases. For example, in cases where LOO-CV scores suggest `Exponential` to be the best choice, convergence plots show the contrary. As another example, in Figure 4 the `PI + Matern32` candidate performs worse than `PI + RBF` candidate although `Matern32` prior has better LOO-CV score than `RBF` prior (see Table S1).

### 4.4 Automatic prior selection

Because the results in Sections 4.2 and 4.3 suggest that different priors fit different datasets, it is natural to consider automatic prior selection. We thus consider the following two new candidate pipelines. They select the prior that maximizes the LOO-CV score on initial design data (all 4 possible priors are considered) and use one of acquisition functions that performed best in previous experiments, namely `PI` and `LogEI`.

We run these independently 64 times on 11 datasets from Section 2.3. The histograms of prior selection frequencies are presented in Figure 5. The plots of convergence on 200 iterations are presented in Supplementary materials, in Figures S4–S6. According to the histograms, frequency of prior selection differs among datasets, however, `RBF` and `Matern52` are clearly the most frequent choices.

**Fig. 5:**
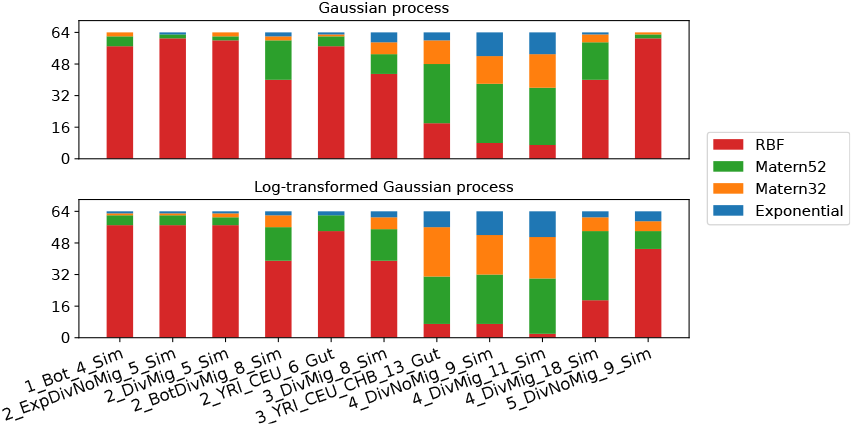
Histograms of automatic prior selection frequencies. For each dataset the top row corresponds to the `PI + Auto` pipeline, while the second row corresponds to the `LogEI + Auto` pipeline.

Upon examining convergence plots it is clear that both new candidates show equal or better performance than the previous champions. For different datasets, especially corresponding to larger population counts, there was no clear winner. For instance, for three datasets `3_DivMig_8_Sim`, `4_DivNoMig_9_Sim` and `4_DivMig_11_Sim` the pipeline `LogEI + Auto` shows better performance than `PI + Auto`. However, for two datasets `3_YRI_CEU_CHB_13_Gut` and `5_DivNoMig_9_Sim` the results turned out the other way around.

### 4.5 Ensembling

Ensembling optimization techniques is a promising tool. By combining (e.g. by simply alternating) several methods it is often able to give a considerably more efficient technique. For example, an ensemble Bayesian optimization technique named Squirrel (Awad *et al*., 2020) was one of the prize winners of the *Black-box Optimization Challenge* in 2020.

Basing on these ideas, we propose yet another candidate pipeline. First, basing on the results of Sections 4.2 and 4.3 and the frequencies of automatic prior choice examined in Section 4.4, we narrow down the set of priors to `RBF` and `Matern52` and the set of acquisitions to `PI` and `LogEI`, deeming them the most efficient. The pipeline starts by chosing the best of the two priors based on the LOO-CV scores over the initial design. Then, on each iteration, one of the two acquisitions is chosen randomly (with equal probability). That is, on different iterations, different acquisition strategies are used. This pipeline is denoted by `Ensemble`.

Comparing `Ensemble` to the previously mentioned best performers by examining convergence plots (Figures S7–S9) shows that the former has similar or better performance. This makes the new `Ensemble` pipeline superior to all other considered candidates and naturally suggests choosing it as the final solution. It shows significant improvements on some datasets, e.g. on `4_DivNoMig_9_Sim` (see Figure 6) and `4_DivMig_11_Sim` (see Figure S4).

**Fig. 6:**
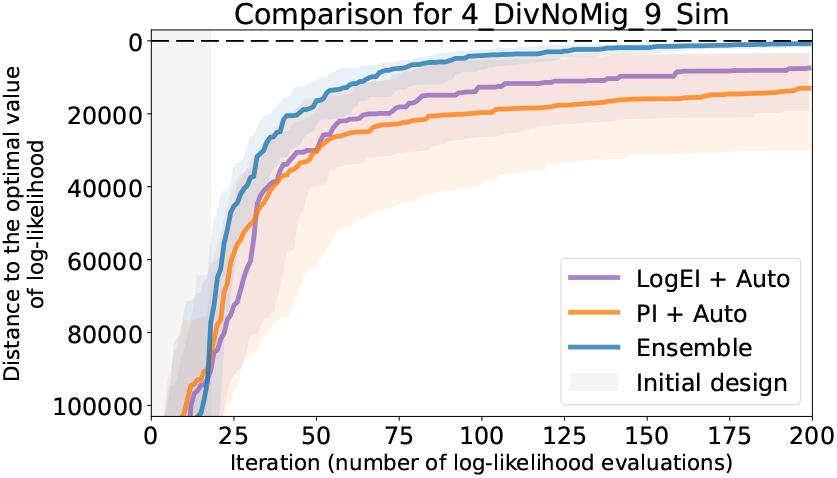
Convergence plots of the previously considered best performers and the new ensemble approach on the `4_DivNoMig_9_Sim` dataset showing superiority of the latter. The meaning of the colored solid lines, shaded regions and the grey area on the left are the same as in Figure 4.

### 4.6 Comparison to the genetic algorithm

Finally, we compare the final champion pipeline `Ensemble` to the genetic algorithm implemented in second version of *GADMA* (Noskova *et al*., 2022). For this, besides the iteration-wise convergence plots used before, we use the wall clock time convergence plots. These are important because, in contrast to the genetic algorithm, Bayesian optimization has tangible computational overhead arising from Gaussian process regression and acquisition function optimization ran on each iteration.

Two additional datasets of modern human populations from Jouganous *et al*. (2017) were included in this evaluation. The convergence plots for thirteen datasets are presented in Figures S10–S14. An example of a wall-clock time convergence plot is presented in Figure 7.

**Fig. 7:**
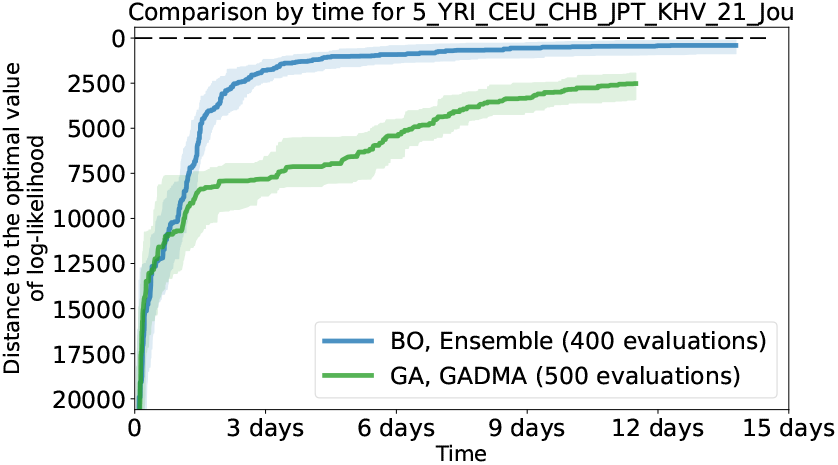
Wall clock time convergence of `Ensemble` pipeline (blue) versus the genetic algorithm implementation from *GADMA* (green) on the `5_YRI_CEU_CHB_JPT_KHV_21_Jou` dataset. The meaning of the colored solid lines and shaded regions are the same as in Figure 4.

The final Bayesian optimization pipeline turns out to be superior to the genetic algorithm on all datasets iteration-wise. However, as mentioned before, Bayesian optimization has overhead in terms of the wall clock time, which makes the wall clock time results different. Here, the more expensive-to-evaluate log-likelihood is, the better Bayesian optimization compares to the genetic algorithm. As it was demonstrated in Figure 3, log-likelihood evaluation time depends on the dataset, mainly on the number of populations therein.

In the end, the genetic algorithm turns out to be superior in terms of wall clock time in cases of one and two populations. In the case of three populations two optimization approaches show comparable results. In the case of four populations Bayesian optimization has faster convergence in the beginning but then ties with the genetic algorithm. For five populations, however, Bayesian optimization turns out to be superior with a margin.

### 4.7 Application to real data

Here, we pay closer attention to the additional datasets by Jouganous *et al*. (2017) that extend the standard out-of-Africa model by Gutenkunst *et al*. (2009) with three populations to the cases of four and five populations.^6^ Jouganous *et al*. (2017) performed demographic inference for these datasets using the *moments*’ simulation engine and Powell’s method with restarts reporting the resulting models and log-likelihood values. Demographic inference for these can bring real insights into the history of humankind and here we show that `Ensemble` is able to find new models for them with higher log-likelihood values then ever previously observed.

The model for four populations has 17 parameters and the model for five has 21: the same 17 ones and four additional ones. To reduce computational time, Jouganous *et al*. (2017) optimized over 17 parameters of the former model, then fixed those and optimized over the remaining four parameters of the second model. We refer to the demographic histories obtained by Jouganous *et al*. (2017) as the *baseline histories*.

First, to infer a history (17 parameters) for the four populations model, we run `Ensemble` 64 times for 400 iterations followed by the BFGS local search until convergence. The results are presented in Table S3. The best result has higher log-likelihood compared to the baseline history. The corresponding history is similar to the baseline but suggests exponential decline of JPT population from 30, 000 to 15, 000 individuals in contrast to the exponential growth from 4, 000 up to 230, 000 individuals in the baseline history. Moreover, the new history suggests a much lower migration rate between CHB and JPT populations. The second best history is much more similar to the baseline but has better value of log-likelihood and lower migration rate between CHB and JPT populations.

After this, we fix the 17 parameters of the five populations model to the same values as in the baseline history, and run `Ensemble` for 200 iterations (followed by the local search) to infer the remaining four parameters. Interestingly, it requires only 16 ± 7 iterations to overrun the baseline log-likelihood. The results are presented in Table S4. The history attaining best log-likelihood predicts migration rate between CHB and KHV populations to be twice as large as in the baseline history. Moreover, the split of CHB population that created KHV population is predicted earlier: 590 generations ago, as opposed to 337 generations in the baseline.

Finally, we use `Ensemble` to infer a demographic history for five populations from scratch, for the full 21 parameter model. We run `Ensemble` for 400 iterations followed by the local search, 64 independent times. The results are presented in Table S4. The best result achieves higher log-likelihood value than both the baseline history and the history inferred by `Ensemble` with 17 parameters fixed. It suggests exponential decline of the JPT population, similar to the best history for four populations. This result is, however, hardly supported by other studies: the population divergence times are estimated to be very early, implying the out-of-Africa event more than a million years ago. The second best model is much better aligned with the contemporary knowledge. The differences in parameters between this model and the baseline model concern mainly YRI and CEU populations. It could be explained by the low number of samples from these two populations in the dataset where allele frequency spectrum was downsized to 5 chromosomes for YRI and CEU populations and down to 30 chromosomes for the other three populations.

## 5 Discussion

We proposed a Bayesian optimization pipeline suitable for demographic inference with expensive log-likelihood evaluations and limited resources. This pipeline was chosen as the single best performer in a series of experiments comparing different Bayesian optimization pipelines. In terms of the number of log-likelihood evaluations before attaining a good result, it was superior to *GADMA*’s genetic algorithm on all datasets. In terms of the wall-clock time, it was superior for optimizing the expensive-to-evaluate log-likelihoods in cases of four and five populations, showing that it can be used to save days and weeks of computations.

The following two paragraphs suggest practical considerations in using the proposed approach, based on its properties and empirical observations.

It is widely presumed that fewer parameters to be optimized make Bayesian optimization more efficient, with 20 being the critical value after which Bayesian optimization is likely to fail (Frazier, 2018). Although in our experiments with up to 21 parameters Bayesian optimization performed well, the setting where 17 parameters were fixed and then only four optimized is noteworthy. There, it took only around 20 log-likelihood evaluations to overrun the best log-likelihood reported in the literature, which is an impressive result. This suggests that iteratively optimizing nested models whenever appropriate might be quite beneficial. We should also note that adjusting the final result of Bayesian optimization by a local search algorithm might be quite important to make the most out of it.

Bayesian optimization is best suited in scenarios with a priori fixed budget, like time and number of computation cores. In our implementation, a user should manually specify the number of iterations (equal to the number of log-likelihood evaluations) that Bayesian optimization should undertake. As noted above, Bayesian optimization could converge quite fast, especially when the parameter count is low. Moreover, because of Bayesian optimization’s own computational overhead (that increases with time), it is not recommended to run it for too many iterations, even if the target function is not expensive. We suggest to keep the number of iterations under 1000, with 200 or 400 being good practical choices.

We note that for up to three populations, *GADMA* allows model free demographic inference by considering more flexible demographic structures (Noskova *et al*., 2020) instead of models. This requires solving optimization problems that involve discrete parameters. Bayesian optimization can be extended to handle these (in fact, the SMAC engine already has this feature) which may in future allow model free demographic inference for four and more populations, if an appropriate alternative to demographic structures is proposed and implemented.

## Footnotes

1 We will use it to model negative log likelihood, which in many cases of interest is indeed positive. This means that Gaussian process regression will be used to predict the logarithm of negative log likelihood.

2 Note that equation (6) is non-rigorous: this form of Bayes’ rule treats the distributions of Gaussian processes as if they were absolutely continuous with respect to a finite dimensional Lebesgue measure, which they are not. The rigorous formalism exists of course, but we do not dwell on this here.

3 Searching for 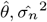 requires solving a linear system and computing a determinant at each iteration of the optimization procedure making it the super-cubic bottleneck in the computational complexity of the regression.

4 https://github.com/noscode/demographic_inference_data.

5 Histogram of joint distribution of derived alleles of individuals.

6 These are exactly the dataset `4_YRI_CEU_CHB_JPT_17_Jou` and the dataset `5_YRI_CEU_CHB_JPT_KHV_21_Jou`, respectively.

