## Supplementary materials for "Bayesian optimization for demographic inference"

[1]Computer Technologies Laboratory, ITMO University, St. Petersburg, 197101, Russia
[2]Department of Computer Science, ETH Zürich, Zürich, 8092, Switzerland
[⋆]Denotes corresponding author

Table S1: Leave-one-out cross validation (LOO-CV) scores for different Gaussian Process priors. The best value for each dataset is in bold.

| | Gaussian process prior | | | |
|---|---|---|---|---|
| Dataset name | `Exponential` | `Matern32` | `Matern52` | `RBF` |
| `1_Bot_4_Sim` | −262.6 | 947.1 | **1320.3** | 1072.1 |
| `2_ExpDivNoMig_5_Sim` | −610.3 | 54.9 | 350.8 | **778.9** |
| `2_DivMig_5_Sim` | −1062.3 | −818.4 | −657.6 | **−373.5** |
| `2_BotDivMig_8_Sim` | **−1514.5** | −1549.0 | −1578.5 | −1689.5 |
| `2_YRI_CEU_6_Gut` | −1118.6 | −921.4 | −788.5 | **−485.0** |
| `3_DivMig_8_Sim` | −1764.5 | −1694.8 | **−1664.3** | −1678.9 |
| `3_YRI_CEU_CHB_13_Gut` | −2027.8 | −1978.7 | −1952.2 | **−1891.4** |
| `4_DivNoMig_9_Sim` | −1532.8 | −1094.8 | **−1082.0** | −1351.5 |
| `4_DivMig_11_Sim` | −1739.9 | −1288.8 | **−1138.7** | −1330.2 |
| `4_DivMig_18_Sim` | −2250.8 | −2222.6 | **−2220.0** | −2235.2 |
| `5_DivNoMig_9_Sim` | −1662.5 | −1149.4 | **−860.1** | −934.1 |

Table S2: Leave-one-out cross validation (LOO-CV) scores for different Gaussian Process priors with log-transformed data. The best value for each dataset is marked in bold.

| Dataset name | Gaussian process prior (log-transformed data) | | | |
|---|---|---|---|---|
| | Exponential | Matern32 | Matern52 | RBF |
| 1_Bot_4_Sim | −200.2 | 681.3 | **947.1** | 830.9 |
| 2_ExpDivNoMig_5_Sim | 192.2 | 863.2 | **972.3** | 441.0 |
| 2_DivMig_5_Sim | −528.7 | 13.8 | 186.1 | **254.8** |
| 2_BotDivMig_8_Sim | −887.0 | **−847.2** | −885.5 | −1039.0 |
| 2_YRI_CEU_6_Gut | −428.7 | 2.6 | **99.9** | 64.1 |
| 3_DivMig_8_Sim | −1147.2 | −890.5 | **−827.0** | −960.6 |
| 3_YRI_CEU_CHB_13_Gut | −1347.0 | **−1251.8** | −1254.5 | −1315.5 |
| 4_DivNoMig_9_Sim | −1277.1 | −750.7 | **−657.3** | −743.6 |
| 4_DivMig_11_Sim | −1180.0 | −845.6 | **−843.1** | −1015.3 |
| 4_DivMig_18_Sim | **−1677.2** | −1681.3 | −1699.2 | −1745.6 |
| 5_DivNoMig_9_Sim | −1132.9 | −403.7 | **37.4** | −3.5 |

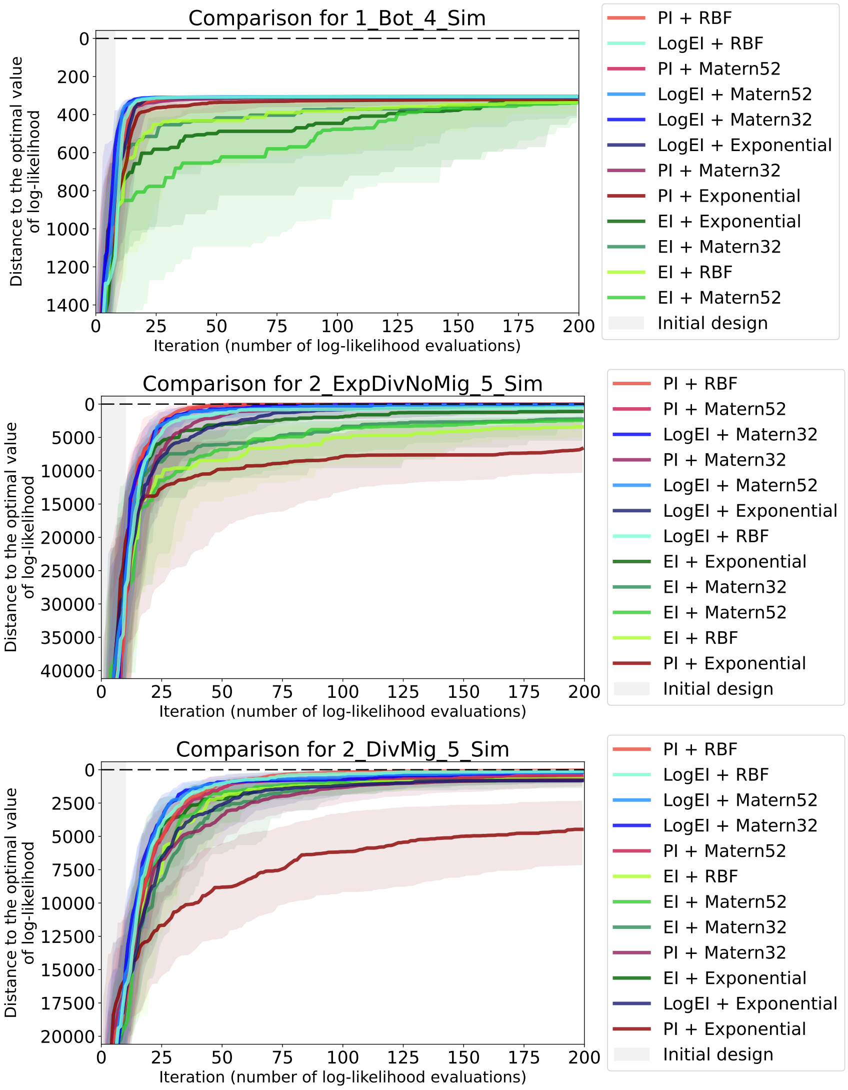


Figure S1: Convergence plots for 12 basic Bayesian optimization pipelines with different acquisition functions and priors. Datasets for **one and two** populations are presented. For each candidate pipeline 64 optimization runs were independently performed. Solid lines of different colors visualize the median of the sample of 64 values on each iteration, while shaded regions visualize ranges between the first and third quartiles. Grey area indicates the initial design, where random search is performed. The labels on legends are sorted according to the median values on the last iteration.

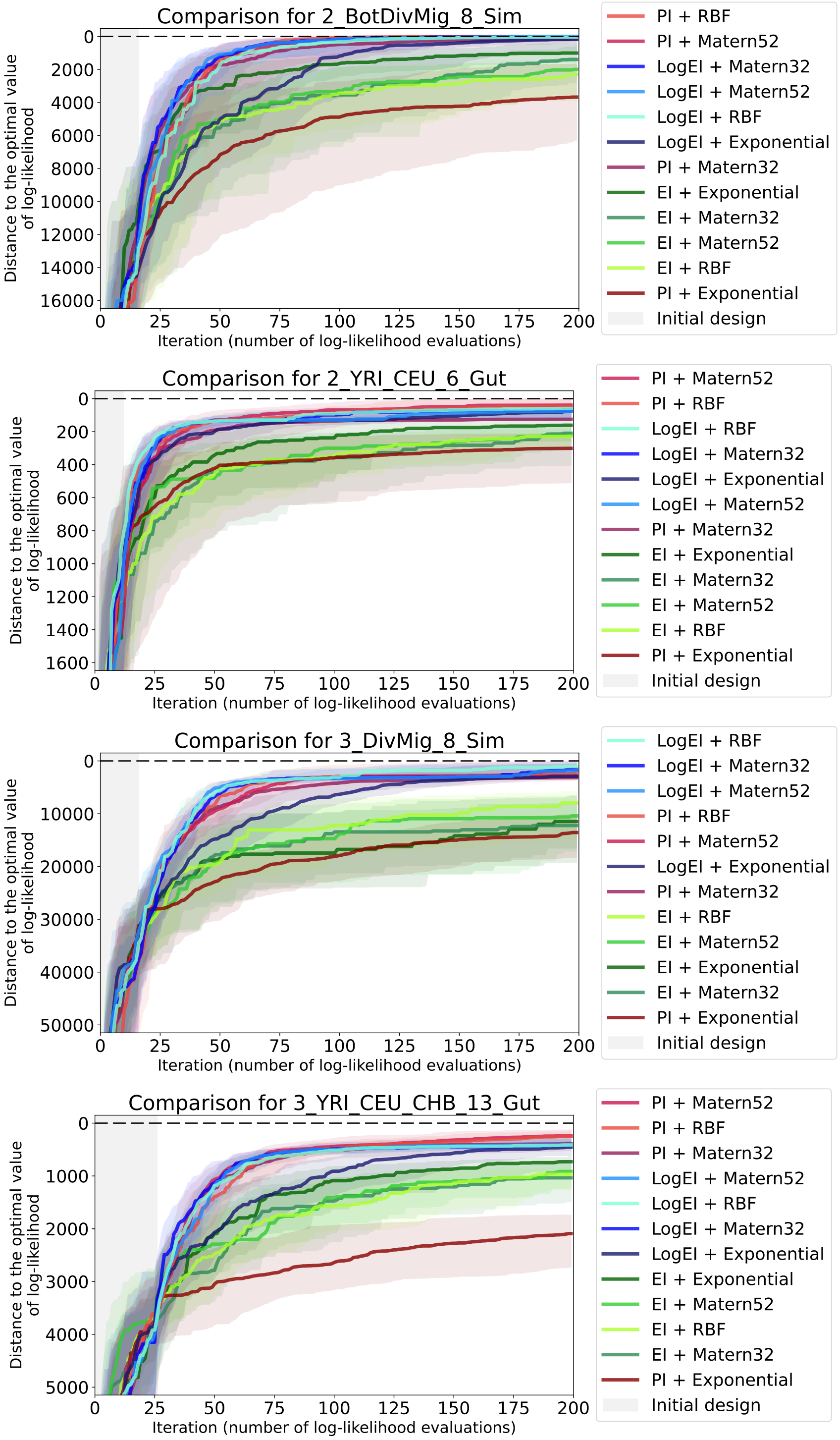


Figure S2: Convergence plots of 12 basic Bayesian optimization pipelines with different acquisition functions and priors. Datasets for **two and three** populations are presented. The labels on legends are sorted according to the median values on the last iteration. The meaning of the colored solid lines, shaded regions and the grey area on the left are the same as in Figure S1.

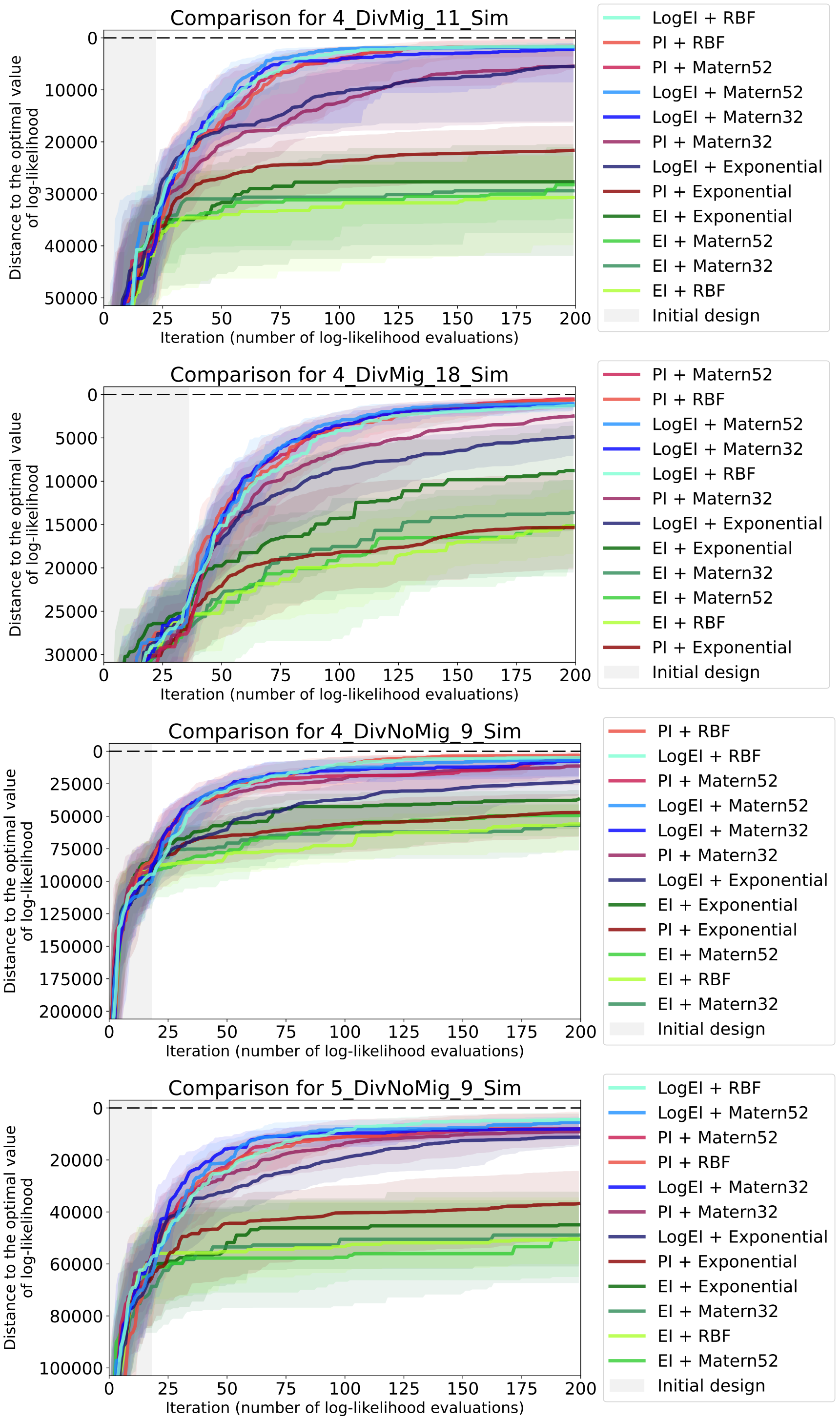


Figure S3: Convergence plots of 12 basic Bayesian optimization pipelines with different acquisition functions and priors. Datasets for **four and five** populations are presented. The labels on legends are sorted according to the median values on the last iteration. The meaning of the colored solid lines, shaded regions and the grey area on the left are the same as in Figure S1.

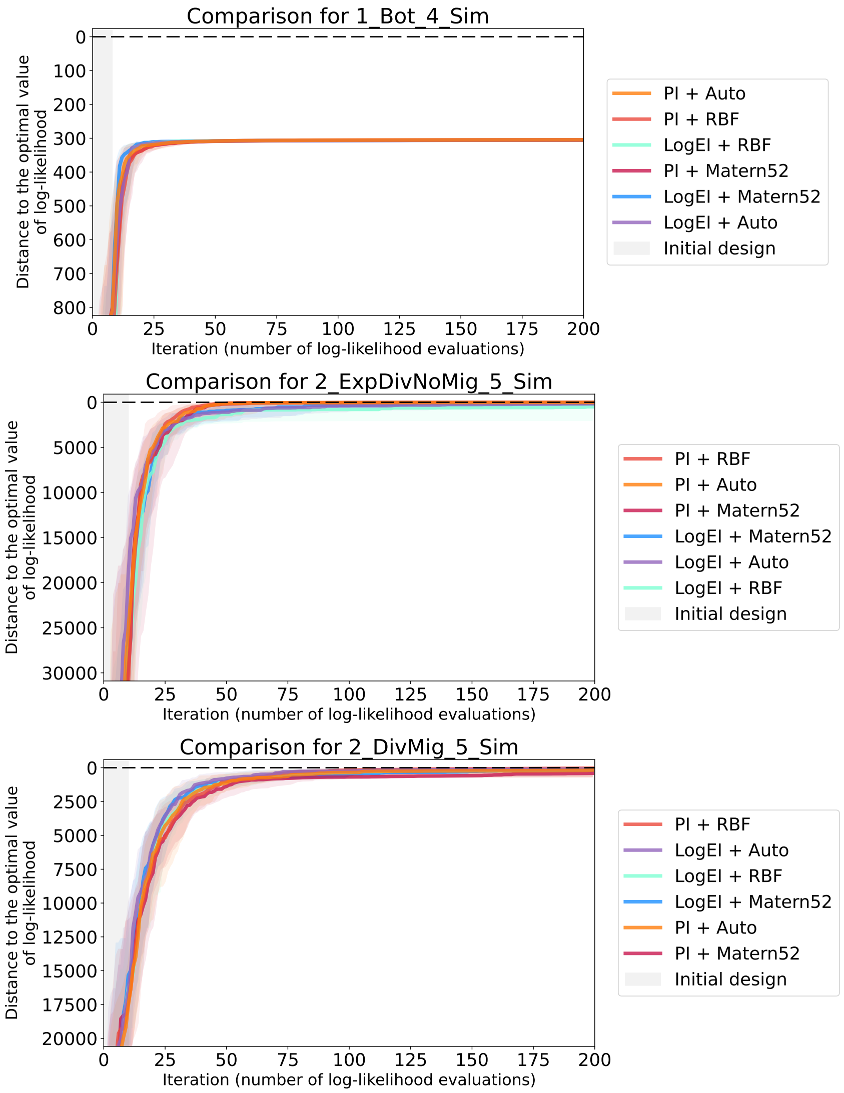


Figure S4: Convergence plots of the previously considered best performers and Bayesian optimization pipelines with automatic prior selection. Datasets for **one and two** populations are presented. For each candidate pipeline 64 optimization runs were independently performed. Solid lines of different colors visualize the median of the sample of 64 values on each iteration, while shaded regions visualize ranges between the first and third quartiles. Grey area indicates the initial design, where random search is performed. The labels on legends are sorted according to the median values on the last iteration.

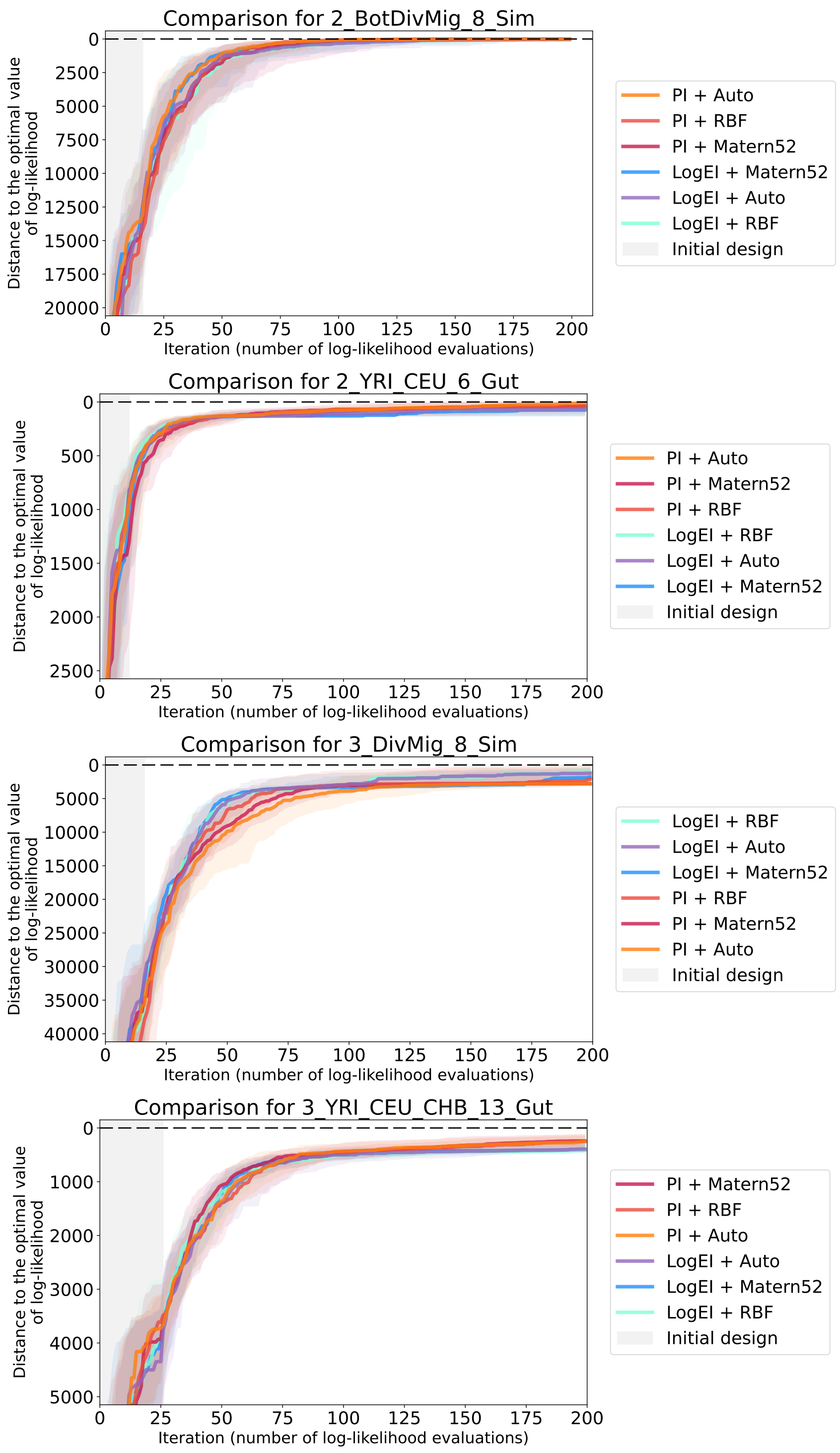


Figure S5: Convergence plots of the previously considered best performers and Bayesian optimization pipelines with automatic prior selection. Datasets for **two and three** populations are presented. The labels on legends are sorted according to the median values on the last iteration. The meaning of the colored solid lines, shaded regions and the grey area on the left are the same as in Figure S4.

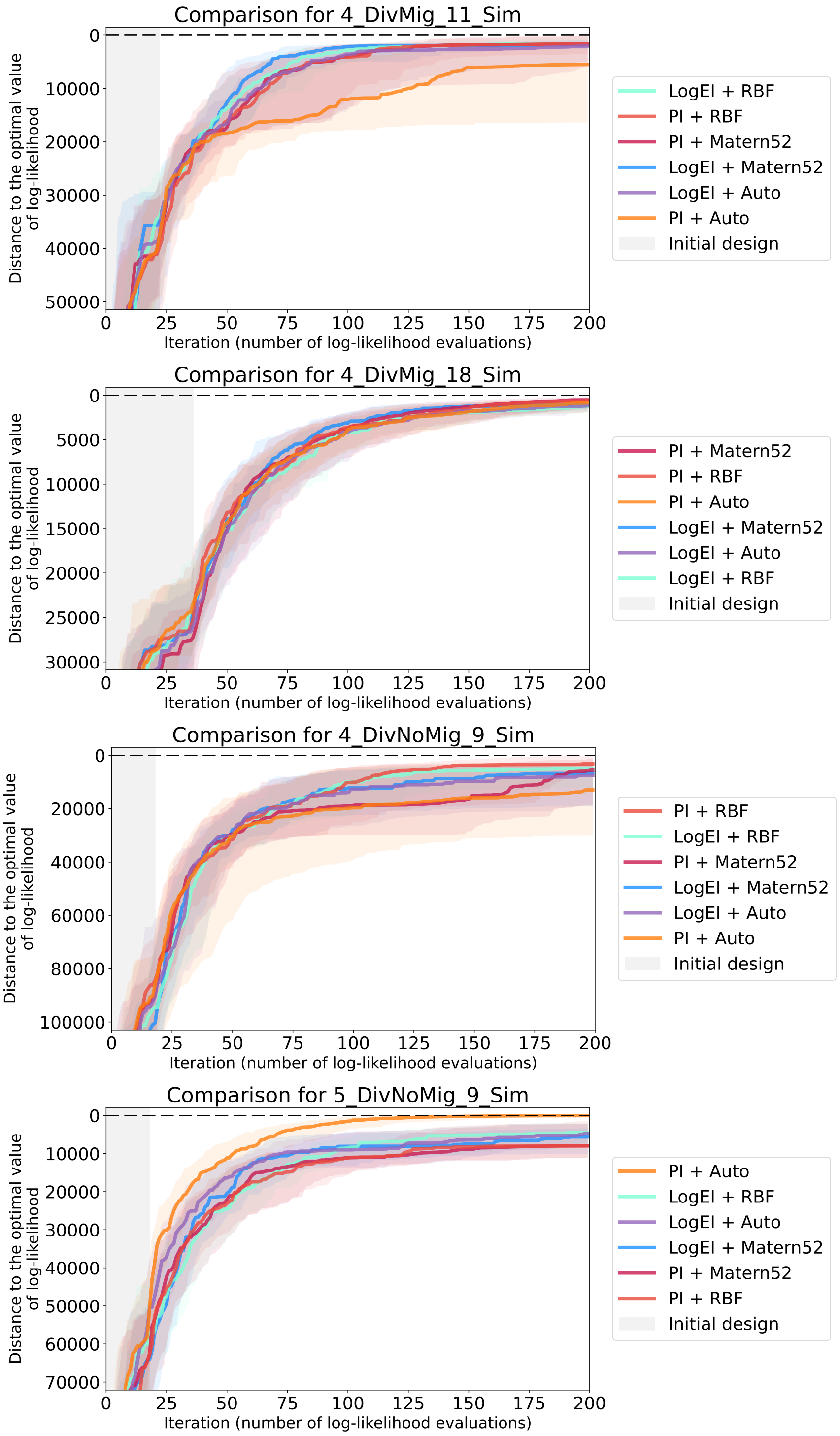


Figure S6: Convergence plots of the previously considered best performers and Bayesian optimization pipelines with automatic prior selection. Datasets for **four and five** populations are presented. The labels on legends are sorted according to the median values on the last iteration. The meaning of the colored solid lines, shaded regions and the grey area on the left are the same as in Figure S4.

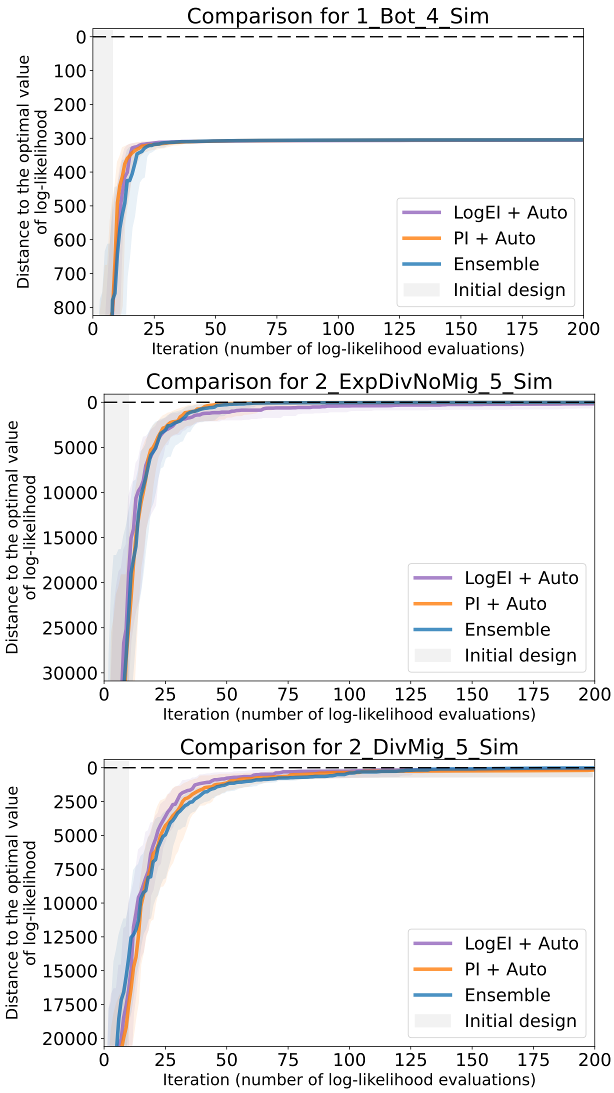


Figure S7: Convergence plots of the Bayesian optimization pipelines with automatic prior selection and the ensemble approach. Datasets for **one and two** populations are presented. For each candidate pipeline 64 optimization runs were independently performed. Solid lines of different colors visualize the median of the sample of 64 values on each iteration, while shaded regions visualize ranges between the first and third quartiles. Grey area indicates the initial design, where random search is performed. The labels on legends are sorted according to the median values on the last iteration.

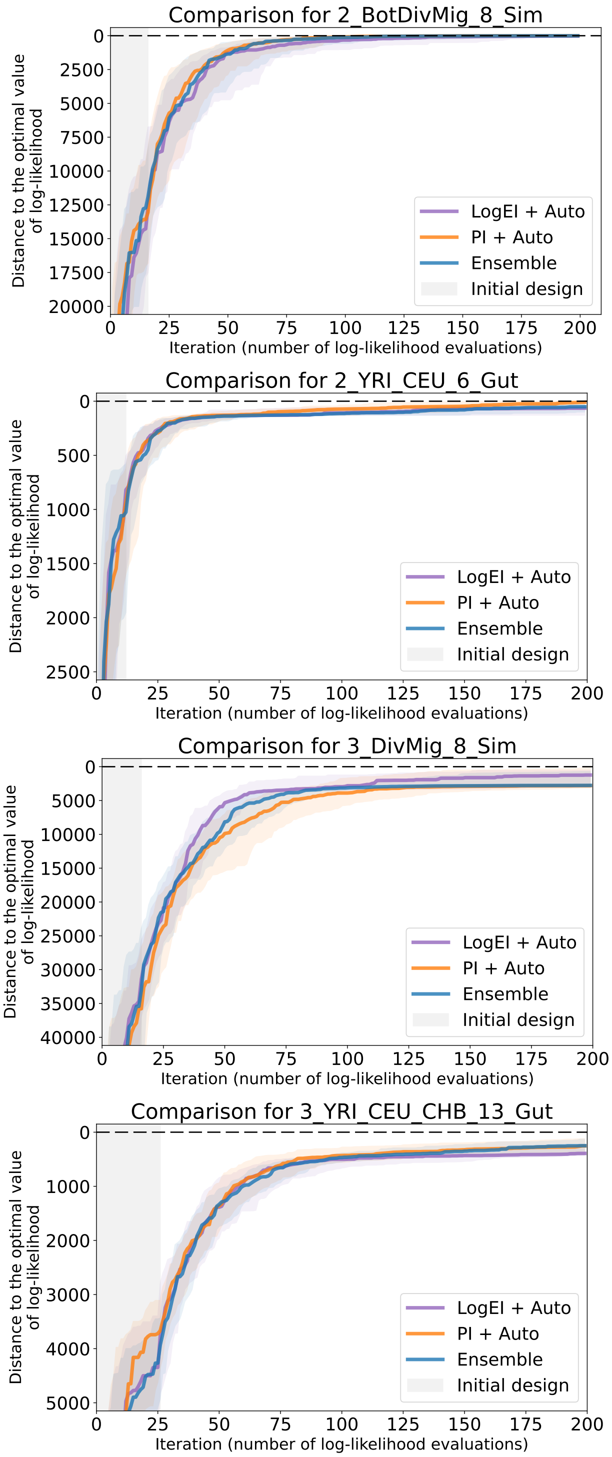


Figure S8: Convergence plots of the Bayesian optimization pipelines with automatic prior selection and the ensemble approach. Datasets for **two and three** populations are presented. The labels on legends are sorted according to the median values on the last iteration. The meaning of the colored solid lines, shaded regions and the grey area on the left are the same as in Figure S7.

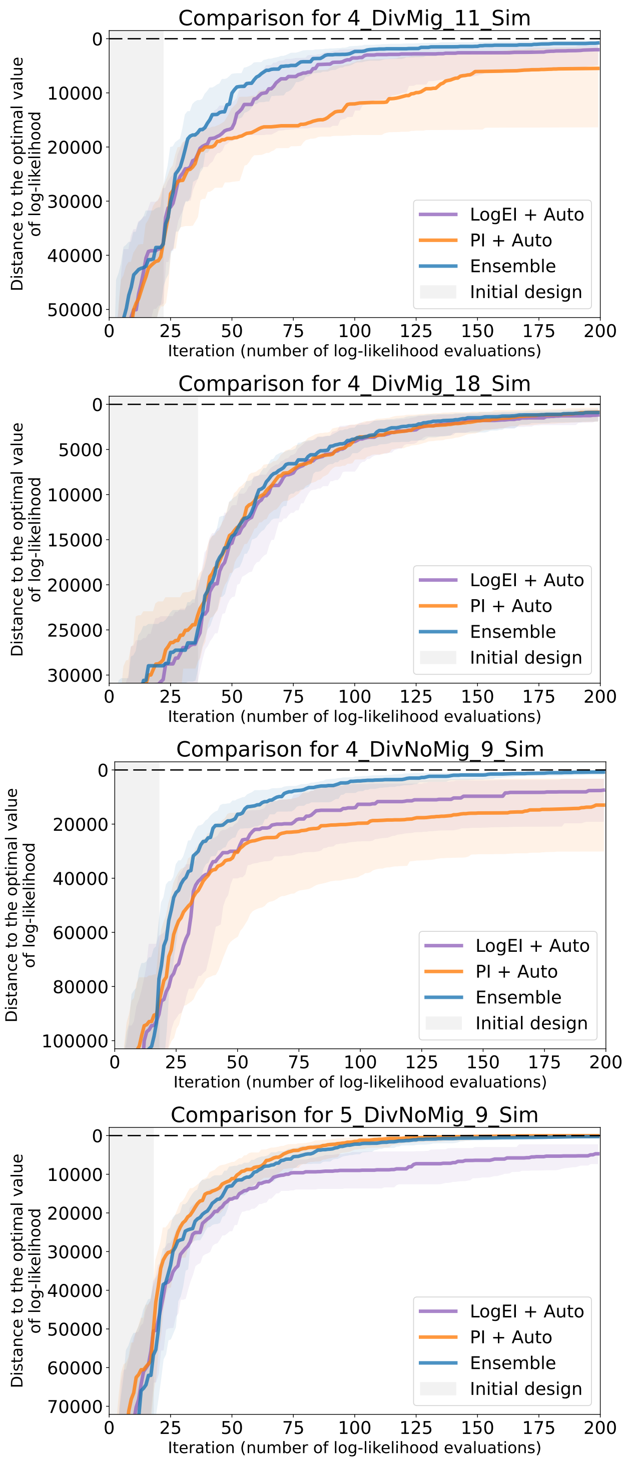


Figure S9: Convergence plots of the Bayesian optimization pipelines with automatic prior selection and the ensemble approach. Datasets for **four and five** populations are presented. The labels on legends are sorted according to the median values on the last iteration. The meaning of the colored solid lines, shaded regions and the grey area on the left are the same as in Figure S7.

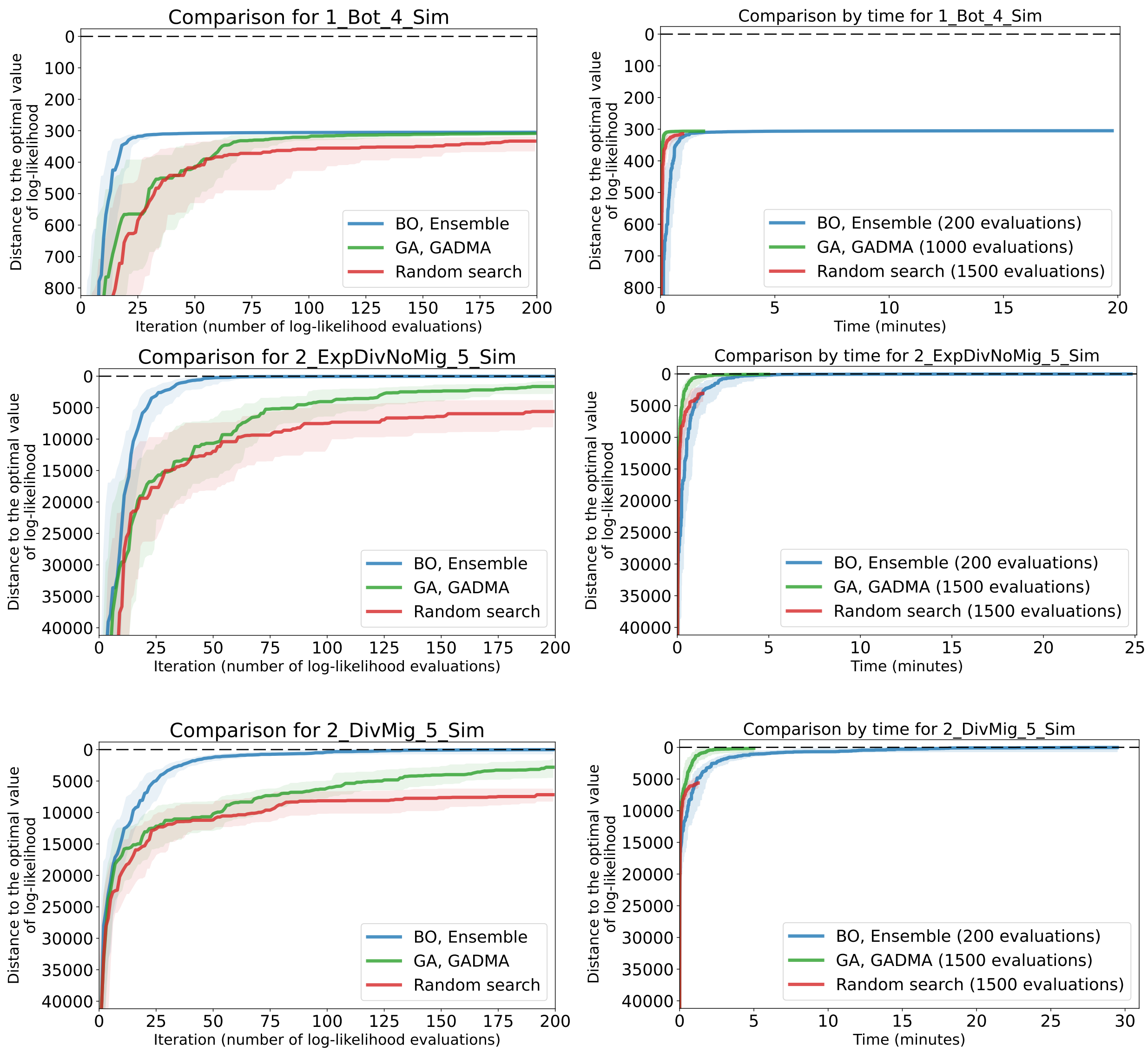


Figure S10: Convergence plots of the Bayesian optimization (BO) ensemble approach, genetic algorithm (GA) and random search. Datasets for **one and two** populations are presented. For each candidate pipeline 64 optimization runs were independently performed. Solid lines of different colors visualize the median of the sample of 64 values on each iteration, while shaded regions visualize ranges between the first and third quartiles. Plots on the left present iteration-wise comparison (200 iterations), plots on the right — wall clock time comparison. Number of evaluations in optimization algorithms used for the wall clock time convergence plots' construction is given in the legends.

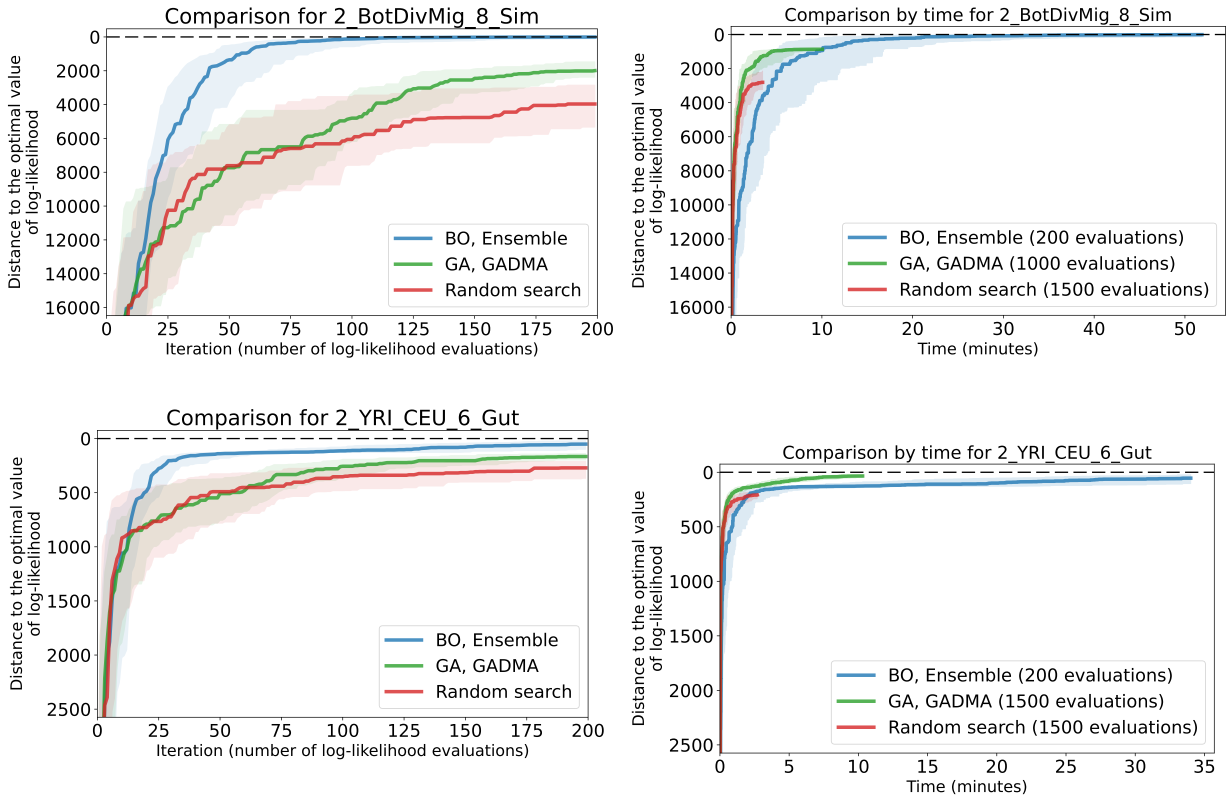


Figure S11: Convergence plots of the Bayesian optimization (BO) ensemble approach, genetic algorithm (GA) and random search. Datasets for **two** populations are presented. Plots on the left present iteration-wise comparison (200 iterations), plots on the right — wall clock time comparison. Number of evaluations in optimization algorithms used for the wall clock time convergence plots' construction is given in the legends. The meaning of the colored solid lines and shaded regions are the same as in Figure S10.

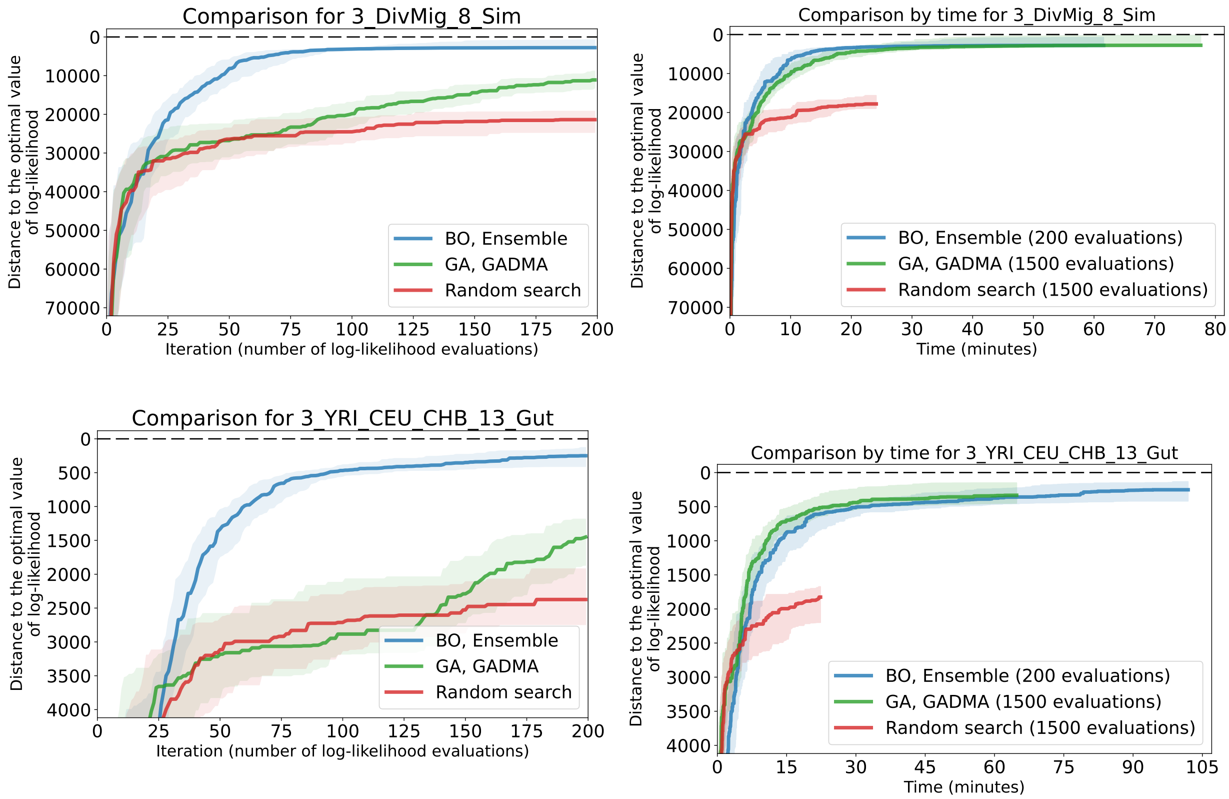


Figure S12: Convergence plots of the Bayesian optimization (BO) ensemble approach, genetic algorithm (GA) and random search. Datasets for **three** populations are presented. Plots on the left present iteration-wise comparison (200 iterations), plots on the right — wall clock time comparison. Number of evaluations in optimization algorithms used for the wall clock time convergence plots' construction is given in the legends. The meaning of the colored solid lines and shaded regions are the same as in Figure S10.

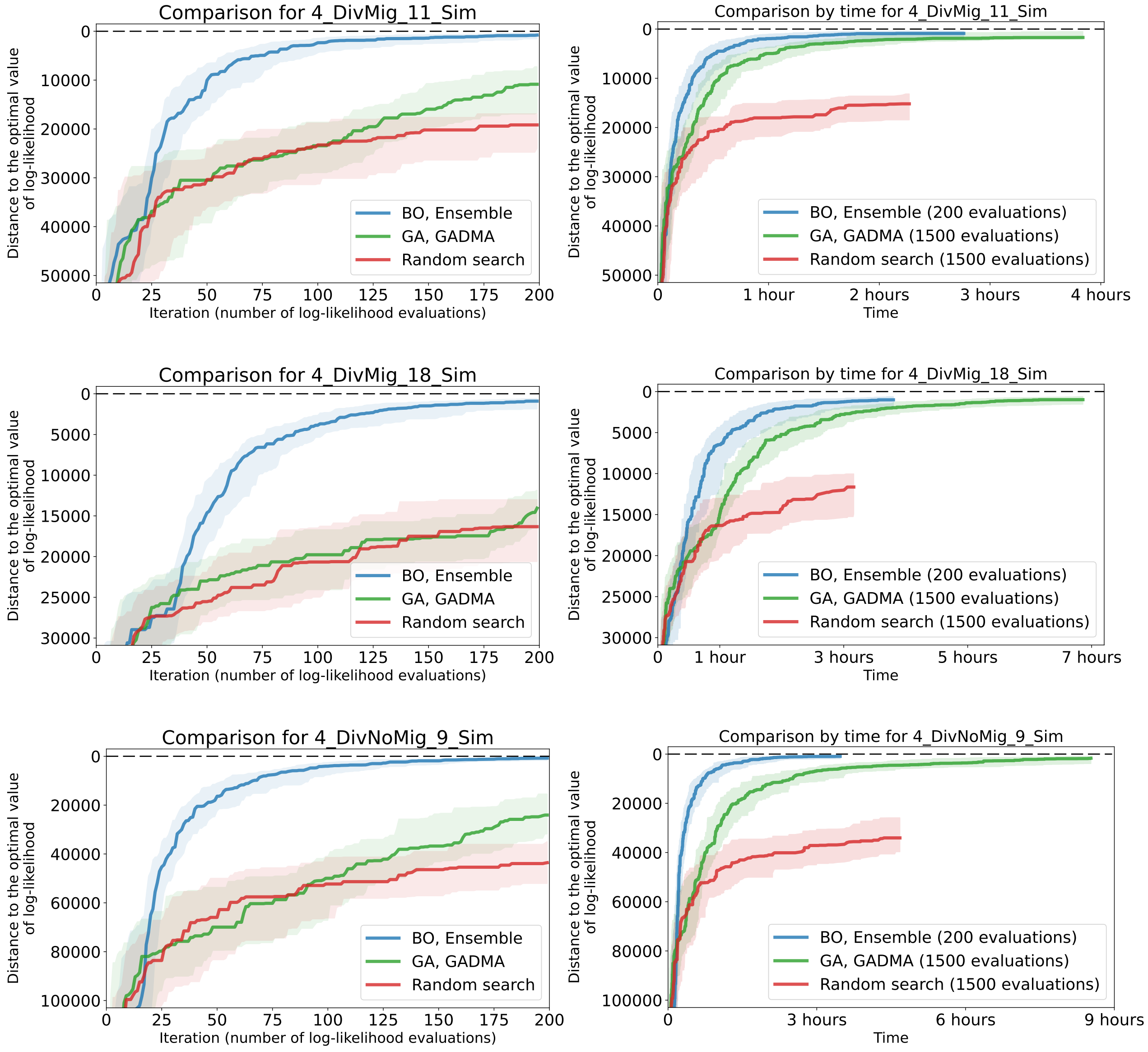


Figure S13: Convergence plots of the Bayesian optimization (BO) ensemble approach, genetic algorithm (GA) and random search. Datasets for **four** populations are presented. Plots on the left present iteration-wise comparison (200 iterations), plots on the right — wall clock time comparison. Number of evaluations in optimization algorithms used for the wall clock time convergence plots' construction is given in the legends. The meaning of the colored solid lines and shaded regions are the same as in Figure S10.

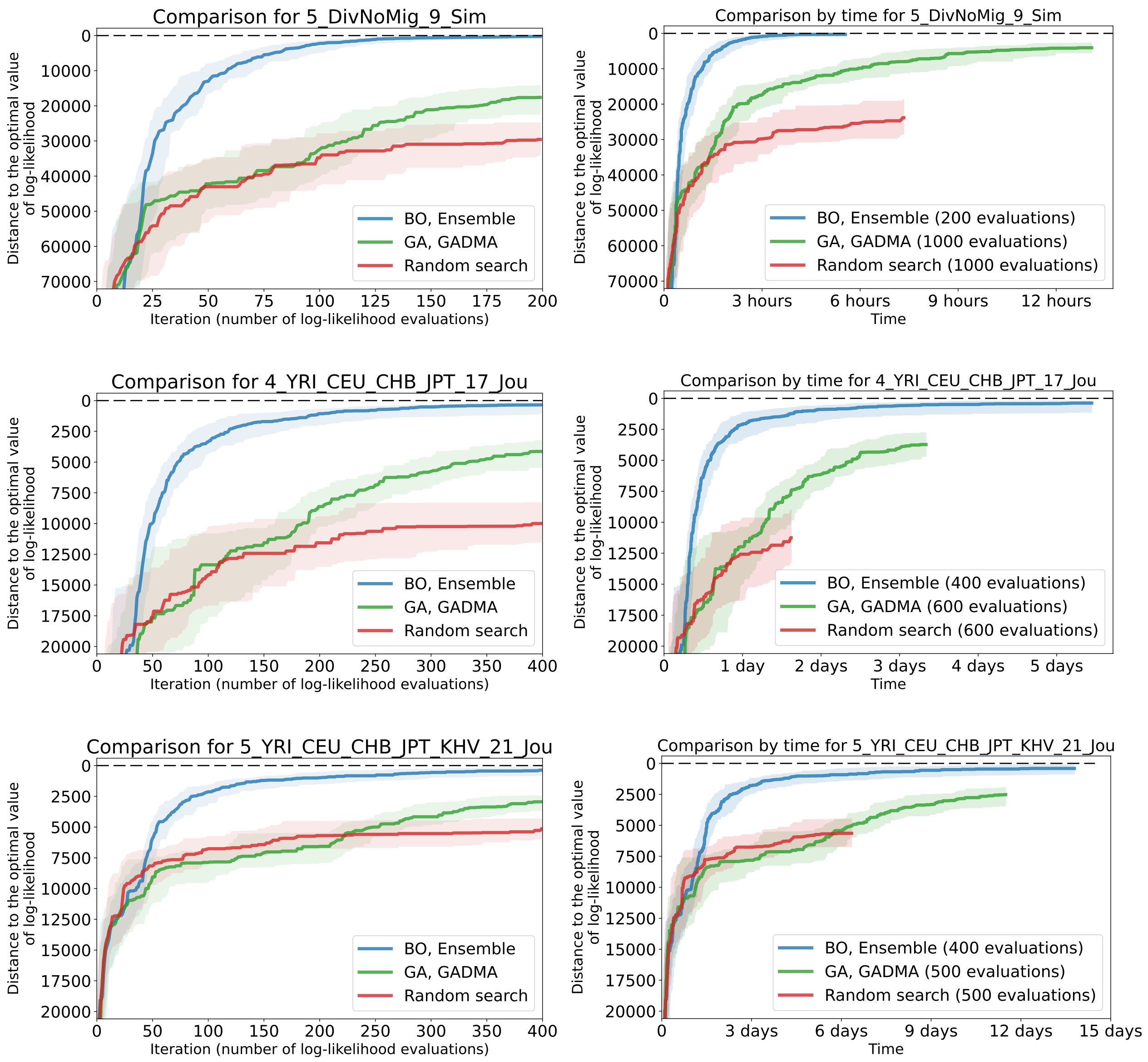


Figure S14: Convergence plots of the Bayesian optimization (BO) ensemble approach, genetic algorithm (GA) and random search. Datasets for **four and five** populations are presented. Plots on the left present iteration-wise comparison (200 or 400 iterations), plots on the right — wall clock time comparison. Number of evaluations in optimization algorithms used for the wall clock time convergence plots' construction is given in the legends. The meaning of the colored solid lines and shaded regions are the same as in Figure S10.

Table S3: Parameters of different demographic histories for four modern human populations (`4_YRI_CEU_CHB_JPT_17_Jou` dataset). Demographic inference is performed in genetic units. The parameters are scaled so that $N_A$ matches the point estimate from Gravel et al. (2011). Two best histories obtained with the Bayesian optimization ensemble approach followed by the BFGS local search are presented as History 1 and History 2.

| | History from Jouganous et al. 2017 | History 1 from BO `Ensemble` | History 2 from BO `Ensemble` |
|---|---|---|---|
| Log-likelihood | −41,852.50 | **−41,837.31** | −41,849.02 |
| Parameters: | | | |
| $N_A$ | 11,293 | 11,293 | 11,293 |
| $N_{AF}$ | 23,721 | 23,178 | 24,081 |
| $N_B$ | 2,831 | 2,520 | 2,613 |
| $N_{EU0}$ | 2,512 | 2,660 | 3,365 |
| $N_{EU}$ | 31,721 | 30,475 | 24,568 |
| $N_{AS0}$ | 1,019 | 1,221 | 1,381 |
| $N_{AS}$ | 62,652 | 44,049 | 41,886 |
| $N_{JP0}$ | 4,384 | 30,224 | 9,507 |
| $N_{JP}$ | 234,113 | 14,517 | 42,678 |
| $m_{AF-B}(\times 10^{-5})$ | 16.8 | 17.77 | 14.81 |
| $m_{AF-EU}(\times 10^{-5})$ | 1.14 | 1.13 | 1.33 |
| $m_{AF-AS}(\times 10^{-5})$ | 0.56 | 0.62 | 0.70 |
| $m_{EU-AS}(\times 10^{-5})$ | 4.75 | 4.78 | 4.37 |
| $m_{CH-JP}(\times 10^{-5})$ | 3.30 | $< 0.01$ | $< 0.01$ |
| $T_{AF}$ (gen.) | 12,310 | 11,164 | 12,042 |
| $T_B$ (gen.) | 4,103 | 3,697 | 3,705 |
| $T_{EU-AS}$ (gen.) | 1,586 | 1,627 | 1,698 |
| $T_{CH-JP}$ (gen.) | 310 | 307 | 297 |

Table S4: Parameters of different demographic histories for five modern human populations. Two models of the demographic history are used: a) with fixed parameters from four populations history and 4 free parameters associated with KHV population, b) the full model with 21 parameters (`4_YRI_CEU_CHB_JPT_KHV_21_Jou` dataset). All parameters are scaled so that $N_A$ matches the point estimate from Gravel et al. (2011). History 1 corresponds to the demographic model with 4 parameters from Jouganous et al. (2017). Values of parameters fixed during inference are underlined. Two best models obtained with the Bayesian optimization ensemble approach followed by the BFGS local search for 21 parameters model are presented as History 2 and History 3.

| | History from Jouganous et al. 2017 | History 1 from BO `Ensemble` | History 2 from BO `Ensemble` | History 3 from BO `Ensemble` |
|---|---|---|---|---|
| Parameters count | 4 | 4 | 21 | 21 |
| Log-likelihood | −34,309.66 | −34,219.41 | **−34,126.20** | −34,155.12 |
| Parameters: | | | | |
| $N_A$ | <u>11,293</u> | <u>11,293</u> | 11,293 | 11,293 |
| $N_{AF}$ | <u>23,721</u> | <u>23,721</u> | 51,347 | 25,645 |
| $N_B$ | <u>2,831</u> | <u>2,831</u> | 6,303 | 2,137 |
| $N_{EU0}$ | <u>2,512</u> | <u>2,512</u> | 5,233 | 17,151 |
| $N_{EU}$ | <u>31,721</u> | <u>31,721</u> | 110,857 | 5,135 |
| $N_{AS0}$ | <u>1,019</u> | <u>1,019</u> | 1,729 | 1,537 |
| $N_{AS}$ | <u>62,652</u> | <u>62,652</u> | 51,046 | 46,196 |
| $N_{KHV0}$ | 2,356 | 5,161 | 1,005 | 7,082 |
| $N_{KHV}$ | 196,813 | 151,576 | 181,310 | 151,132 |
| $N_{JP0}$ | <u>4,384</u> | <u>4,384</u> | 113,089 | 1,326 |
| $N_{JP}$ | <u>234,113</u> | <u>234,113</u> | 27,185 | 246,308 |
| $m_{AF-B}(\times 10^{-5})$ | <u>16.8</u> | <u>16.8</u> | 10.82 | 12.18 |
| $m_{AF-EU}(\times 10^{-5})$ | <u>1.14</u> | <u>1.14</u> | $< 0.01$ | $< 0.01$ |
| $m_{AF-AS}(\times 10^{-5})$ | <u>0.56</u> | <u>0.56</u> | $< 0.01$ | 0.48 |
| $m_{EU-AS}(\times 10^{-5})$ | <u>4.75</u> | <u>4.75</u> | 2.67 | 2.45 |
| $m_{CH-KHV}(\times 10^{-5})$ | 21.30 | 44.27 | 43.77 | 16.80 |
| $m_{CH-JP}(\times 10^{-5})$ | <u>3.30</u> | <u>3.30</u> | 28.33 | 14.44 |
| $T_{AF}$ (gen.) | <u>12,310</u> | <u>12,310</u> | 149,862 | 6,727 |
| $T_B$ (gen.) | <u>4,103</u> | <u>4,103</u> | 84,109 | 3,009 |
| $T_{EU-AS}$ (gen.) | <u>1,586</u> | <u>1,586</u> | 3,818 | 1,706 |
| $T_{CH-KHV}$ (gen.) | 337 | 590 | 3,802 | 411 |
| $T_{CH-JP}$ (gen.) | <u>310</u> | <u>310</u> | 964 | 188 |
